# Strong STMP-Crosslinked Lignin/Chitosan Hydrogel Films with Enhanced Aqueous Stability and Bioactivity for Active Food Packaging

**DOI:** 10.1101/2025.11.11.687780

**Authors:** Sumona Garg, Althuri Avanthi

## Abstract

Plastic pollution has intensified globally due to the widespread use of non-biodegradable packaging materials. Conventional passive plastics used in food packaging lack adaptive functionality limiting their preservation efficiency. Biopolymer-based hydrogels offer promise due to their biodegradability, film-forming ability, and intrinsic bioactivity. However, their instability in aqueous environments constrains their application in high-moisture foods such as meat. Lignin extracted from sugarcane tops via acid–alkali treatment and chitosan known for its film-forming and antimicrobial properties were combined to enhance the antioxidant, antimicrobial, UV-blocking, and mechanical performance of the hydrogel. The films were cured at 25 °C under 80% relative humidity, eliminating the high temperature pre-oven drying required in conventional methods. Further, a stable lignin/chitosan hydrogel film was developed using sodium trimetaphosphate (STMP) as a crosslinker and glycerol as a plasticizer. The films treated with 1% (w/v) STMP for 15 min showed optimal performance, exhibiting 89.89% radical scavenging activity, complete UV shielding, antimicrobial activity against *E. coli*, and tensile strength of 2364.79 MPa with 31.49% elongation at break. The film retained its integrity in food simulants namely ethanol (10% v/v), acetic acid (3% v/v), and distilled water. In chicken breast packaging trials, the CL-0.8/3Cht film-maintained pH, moisture, and protein stability and significantly reduced microbial load during 5 days of storage. These findings establish the CL-0.8Lig/3Cht hydrogel film with agri-residue derived lignin as a robust, multifunctional, and biodegradable active packaging material for moisture-rich food systems. These findings support waste valorisation advancing cleaner and sustainable food preservation practices.

## 1.0 Introduction

The growing environmental burden associated with petrochemical-based plastics has intensified global research in bio-based and biodegradable alternatives for sustainable food packaging applications. According to recent estimates, plastic packaging accounts for over 40% of global plastic waste, with nearly one-third attributed to food packaging applications (Geyer et al., 2017)(Ncube et al., 2021). While various bio-derived polymers have been explored, most remain limited by trade-offs in functionality, scalability, and biodegradability. Among these, chitosan is a polycationic polysaccharide sourced from crustacean exoskeletons has garnered significant interest due to its inherent antimicrobial activity, film-forming capacity, and biodegradability. However, its practical utility in food packaging, especially for moisture-rich perishable products such as raw meat, remains severely constrained. This is largely due to its poor mechanical integrity and high-water sensitivity under humid or aqueous environments (Mao et al., 2019). These hydrophilic limitations significantly restrict the deployment of these films in real-world food packaging applications.

Although several studies attempted to improve film forming abilities of chitosan through blending, plasticization, or crosslinking, most fabrication methods are relied on synthetic or potentially toxic additives. These approaches have not effectively addressed the dual challenge of retaining mechanical strength and functional stability under varying humidity conditions and in food simulants (such as water, ethanol, and acetic acid) (Mao et al., 2019) (Pei et al., 2025)(Orcutt et al., 2025). Furthermore, the sustainable development goals (SDGs) proposed by United Nations General Assembly (UNGA) stresses on valorisation of underutilized biomass waste streams that aligns with principles of green chemistry (use of renewable feedstocks) and circular economy.

The present study focuses on lignin, the second-most abundant aromatic biopolymer in nature and a largely under-utilised resource in materials development. Despite its rich polyphenolic framework and inherent antioxidant capacity, lignin is routinely treated as waste in lignocellulosic biorefineries such as pulp & paper mills and second[generation biomass conversion plants (Branco et al., 2018). Various forms of lignin, including kraft, soda and organosolv, have been explored for materials applications, yet their heterogeneity and poor solubility often impede film formation. The source and extraction route of lignin markedly influence its reactivity, compatibility and composite performance. Notably, lignin derived from grass family tends to present higher phenolic hydroxyl content than lignin from dicots, rendering it more reactive and favourable for incorporation into biopolymeric composites (Ralph et al., 1998) (Higuchi et al., 1967). In this study, lignin is sourced from sugarcane tops (SCT) and an overlooked agricultural byproduct produced approximately 6–8 tons per hectare of sugarcane produce, (Singh et al., 2008) (Chandel et al., 2012) thus contributing significantly to the biomass waste in major sugarcane-growing regions. Unutilized SCT is an environmental burden and to mitigate its accumulation, farmers often resort to open field burning or landfilling, both of which contribute substantially to greenhouse gas emissions and environmental pollution.

The objective of this study is to transforms agri-waste into high-performance, biodegradable packaging material. Despite rich functional chemistry of lignin, its direct incorporation into chitosan matrices typically leads to poor film homogeneity and instability in aqueous environments due to weak physical and chemical linkages. In this study, this bottleneck was addressed by employing sodium trimetaphosphate (STMP), a food-grade crosslinker known to form phosphoamino bridges with amino groups of chitosan and has a potential to form phosphoester bonds with hydroxyl groups of lignin. The introduction of STMP can enhance mechanical strength and water stability of the lignin/chitosan matrix, achieving durability under prolonged exposure to moisture which is a critical requirement for meat packaging. Further to minimize brittleness of the crosslinked films, glycerol was introduced as a natural plasticizer. Unlike synthetic alternatives, glycerol is GRAS (Generally Recognized as Safe) by Food and Drug administration and can offer a favourable balance between flexibility and tensile performance.

A comprehensive formulation-to-function investigation of a lignin/chitosan hydrogel film structurally engineered via STMP crosslinking and glycerol plasticization is carried out in this study. Designed explicitly for high-performance food packaging, the film was systematically evaluated for key physicochemical attributes including tensile strength, antioxidant, barrier properties, thermal resilience, and aqueous swelling behaviour under storage conditions. Furthermore, real-time packaging trials were conducted on raw chicken breast stored at 25 °C, an ambient temperature reflective of low-resource environments where cold chain infrastructure is often lacking. These indices provided quantifiable evidence of the film’s effectiveness in retaining freshness, slowing oxidative spoilage, and extending shelf life of chicken breast. Hence, as the packaging industry pivots toward greener technologies, this synthesised bioactive film could play a crucial role in reducing reliance on plastics, minimizing food waste, and promoting circular economy models.

## 2.0 Materials and Methods

### 2.1 Materials

Sugarcane tops (variety VSI12121) were collected from agricultural fields in Kandi, Sangareddy, Telangana, India. The biomass was oven-dried at 60 °C and sieved to obtain particle sizes of 0.25 mm. Low molecular weight chitosan (∼90% degree of deacetylation), lactic acid, Folin–Ciocalteu reagent, bovine serum albumin (BSA), sodium hydroxide, de Man, Rogosa, and Sharpe (MRS) broth, nutrient broth, and agar media were sourced from Sisco Research Laboratories Pvt. Ltd. (SRL), India. Sodium carbonate was obtained from Pure Synth (India), copper(II) sulfate pentahydrate from Honeywell (USA), and potassium sodium tartrate from Chemsynth (India). Sulfuric acid, sodium trimetaphosphate (STMP), and hydrochloric acid were supplied by SD Fine Chemicals and Avra Synthesis Pvt. Ltd. *Escherichia coli* DH5α was donated from chromosomal dynamics research lab in the Department of Biotechnology, IIT Hyderabad.

### 2.2 Preparation of lignin/chitosan hydrogel films

Hydrogel films were prepared with and without lignin as described previously (Garg and Avanthi, 2025). Briefly, 2% (w/v) chitosan in 1% (v/v) lactic acid (150 mL) was adjusted to pH 6–6.5. Lignin extracted with 1% (v/v) H[SO[and 3% (w/v) NaOH was dissolved in 70% ethanol at 60°C for 24 h. Lignin solution (1% w/v) of 80 mL was added to the chitosan solution and stirred at 70°C until reduced to 30% volume. About 75–80 mL of the mixture was casted on Teflon sheet (20 cm× 12 cm), air-dried, and peeled to obtain 0.8Lig/3Cht control film.

### 2.3 Crosslinking with STMP

The 0.8Lig/3Cht films were immersed in STMP solutions (1%, 1.5%, and 2% w/v) for 15, 30, 45, 60, and 75 min, blotted to remove excess solution. The films were again dipped in 50% (v/v) glycerol solution for 5mins followed by blotting. The films were cured overnight at 25°C and 80% RH to complete crosslinking. The resulting films were designated CL-0.8Lig/3Cht.

### 2.4 Characterization

#### 2.4.1 Swelling behaviour analysis

Swelling capacity was measured by immersing pre-weighed dry films (W[) in 10 mL distilled water at room temperature. After set intervals, films were blotted to remove surface water and reweighed (W[). The swelling ratio was then calculated using equation 1.

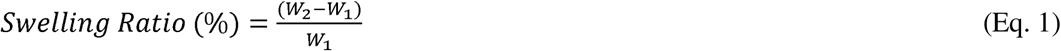

Where W&#x25A1; is the initial dry weight of the film and W&#x25A1; is the weight of the swollen film after immersion (Wu et al., 2019).

#### 2.4.2 Water vapour transmission rate (WVTR) and Radical scavenging activity (RSA)

WVTR was measured gravimetrically by sealing a 150 mL flask containing 10 mL water with a 3 cm × 3 cm film with parafilm. The flask was weighed before (W&#x25A1;) and after 24 h at 40 ± 2°C (W&#x25A1;) (Garg and Avanthi, 2025), and WVTR was calculated using equation 2.

Radical scavenging activity (RSA) was determined by incubating 200 mg of film in 10 mL of 95% (v/v) ethanol at 50°C for 3 h followed by mixing 1.5 mL of this solution with 1.5 mL 0.06 mM DPPH. The reaction mixture was incubated at 30 min in the dark, and the absorbance was taken at 517 nm to calculate RSA using equation 3 (Garg and Avanthi, 2025).

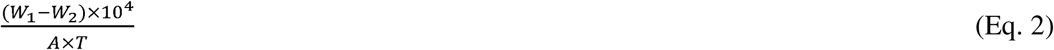

Where t is the time (24 h) and A is the area of the bottle mouth (mm²)

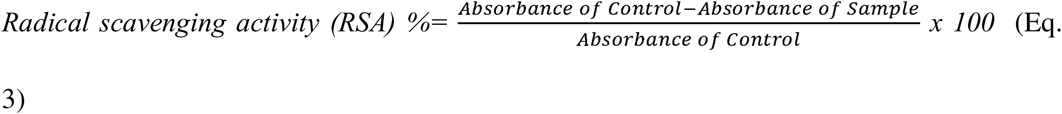

#### 2.4.3 Antimicrobial activity

The antimicrobial properties of the hydrogels were evaluated against *E. coli* DH5α using the disk diffusion assay. Circular hydrogel discs (0.6 mm in diameter) were sterilized by exposure to ultraviolet (UV) light for 15 minutes. Each sterilized disc was then carefully placed on agar plates pre-inoculated with 10 µL of bacterial broth culture (optical density ≈ 0.7). The plates were incubated at 25 °C for 24 hours. Following incubation, the diameters of the inhibition zones around the hydrogel discs were measured to determine antimicrobial effectiveness.

#### 2.4.4 Contact angle

Surface wettability of 0.8Lig/3Cht and CL-0.8Lig/3Cht films was measured using a KRÜSS Drop Shape Analyzer. A 4 µL water droplet was placed on a 2 cm × 2 cm film samples, and the contact angle was determined via the sessile drop method using built-in software(Banerjee et al., 2025).

#### 2.4.5 UV blocking property

The UV-blocking efficiency of CL-0.8Lig/3Cht was measured using a Lambda 365 UV–Vis spectrophotometer over 200–800 nm, with air as the baseline and the results were compared with 0.8Lig/3Cht hydrogels film (Garg and Avanthi, 2025). Samples were cut to fit quartz cuvettes. The Sun Protection Factor (SPF) was calculated from absorbance values in the UV-B range (280–320 nm) using equation 4

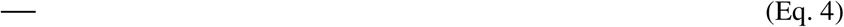

where, FC represents a constant equal to 10, EE(λ) is the erythemal effectiveness, I(λ) denote the solar intensity at a specific wavelength, and Abs(λ) is the absorbance of the sample at that wavelength (Almeida et al., 2019).

Absorbance values were recorded across the 290–320 nm range at 5 nm intervals, and this value was substituted in equation 5.

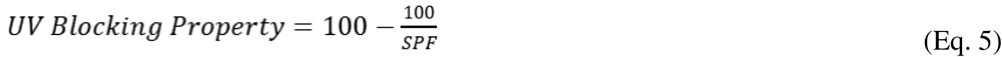

#### 2.4.6 Mechanical testing and x-ray photoelectron spectroscopy (XPS)

Mechanical properties of CL-0.8Lig/3Cht films (20 mm× 20 mm) were measured using an Instron UTM (ASTM D638) and compared with 0.8Lig/3Cht films (Garg and Avanthi, 2025)to evaluated tensile strength, elongation, and Young’s modulus at 10 mm/min. Surface chemistry was analyzed by XPS (AXIS Supra, Kratos Analytical, UK) using Al Kα radiation (1486.6 eV) with spectra acquired at 1 mm spot size, 40 eV pass energy that was calibrated to C 1s 284.6 eV, and processed using ESCApe software.

#### 2.4.7 Thermogravimetric analysis (TGA)

The thermal stability of the films was evaluated using TGA (TA Instruments SDT Q600). Sample of 5 mg was heated from 30°C to 800°C at 10°C/min under nitrogen, and weight loss profiles were recorded to assess degradation behaviour and residual mass. The results were compared with the control film 0.8Lig/3Cht(Garg and Avanthi, 2025)

#### 2.4.8 Differential scanning calorimetry (DSC)

The thermal behaviour of the 3Cht, 0.8Lig/3Cht, and CL-0.8Lig/3Cht was analyzed using DSC (TA Instruments Q2000) under nitrogen atmosphere. Sample of 1mg was sealed in aluminium pans and heated from 50°C to 420°C at 5°C/min (Rodrigues et al., 2020). Limiting oxygen index (LOI) of the films was calculated using Equation 6 to evaluate their flammability characteristics.

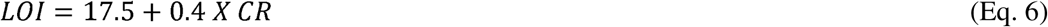

where CR represents the char residue (%) of the films measured at 800 °C (Brenner et al., 2020).

The ignition point and self-extinguishing behaviour of the chitosan (control) and CL-0.8Lig/3Cht (test) films were evaluated by exposing them to a direct flame by a lighter. Ignition and self-extinguishment times were recorded. Sequential images from the video were captured to document combustion progression and structural changes.

### 2.4.9 Morphological analysis

The surface morphology of the films was examined using focused ion beam scanning electron microscopy (FIB-SEM, JEOL JIB-4700). Both air and matrix side of 0.8Lig/3Cht and CL-0.8Lig/3Cht (with and without glycerol) films were gold-coated for 30 s before imaging to ensure conductivity. The samples were designated as M-NC (matrix side of non-crosslinked film), A-NC (air side of non-crosslinked film), M-CL-G (matrix side of crosslinked film with glycerol), A-CL-G (air side of crosslinked film with glycerol), M-CL (matrix side of crosslinked film without glycerol), and A-CL (air side of crosslinked film without glycerol). High-resolution SEM was used to assess surface topography and uniformity and energy dispersive x-ray spectroscopy (EDS) at 15 kV to confirm the elemental composition across the film surface (Yu et al., 2021).

### 2.4.10 Anti-freezing and Anti-wrinkle test

Anti-freezing and anti-wrinkle properties were assessed visually and mechanically. For the anti-freezing test, 3 cm × 1 cm films were stored at −20°C and 4°C, then twisted with forceps to evaluate flexibility and photographed for structural changes. For the anti-wrinkle test, 5 cm × 5 cm films were fully crumpled, rested 10 min at room temperature, and unfolded to observe recovery to their original smooth form.

#### 2.4.11 Soil burial test

Biodegradability was assessed using the soil burial method at 25 ± 2 °C. Pre-weighed film samples 2 cm × 2 cm were buried 5 cm deep in 50 g of red loam soil and kept moist with periodic distilled water sprays. At set intervals (10–90 days), samples were retrieved, and weighed to calculate percentage weight loss using equation 7-

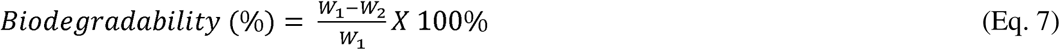

where, W_1_and W_2_ are the initial and remaining dry weights of the film, respectively (CHAISUWAN et al., 2023)

### 2.5 Evaluation of chicken breast preservation performance

#### 2.5.1 Compositional analysis of chicken breast

Ash and moisture content of freshly slaughtered chicken breast samples was determined using muffle furnace at 550 °C for 6 h and weight difference between initial and oven-dried samples at 105 °C, respectively. Lipid and protein content was determined by Bligh and Dyer and Bradford method, respectively.

#### 2.5.2 pH test, weight loss (%), total microbial count (TMC), malondialdehyde (MDA), and soluble protein content

Physicochemical (pH, weight loss %, and total microbial count) of chicken breast samples were conducted following the Manual of Methods of Analysis of Food–Meat and Meat Products*(FSSAI, 2012)*. Lipid oxidation and soluble protein content was evaluated by measuring malondialdehyde (MDA) through the TBA assay (Li et al., 2011) and modified Lowry method (“Quantitation of Total Protein Content in Some Common Edible Food Sources by Lowry Protein Assay,” 2020)

### 2.6 Statistical analysis

All experiments in this study were conducted in triplicates, and the results are presented as mean ± standard deviation. Statistical significance was assessed with p-values < 0.05 considered indicative of significant differences.

## 3.0 Results and Discussions

### 3.1 Concentration-dependent effects of STMP on network stability in water

To assess the effect of crosslinking density on hydrogel performance, chitosan–lignin hydrogels were crosslinked with 0.5%, 1%, and 1.5% (w/v) STMP and immersed in water for 7 days (Fig. 1A). As shown in Fig. 1B, 0.8Lig/3Cht hydrogels disintegrated within 5 s, confirming the necessity of crosslinking. The highest initial swelling was showed with 0.5% STMP (33.20 ± 12.85% on Day 2) but disintegrated by Day 7, indicating inadequate crosslinking. In contrast, the 1% (w/v) STMP hydrogels exhibited lower swelling (16.99 ± 2.39% on Day 2; 12.97 ± 1.28% by Day 3) and remained stable through Day 7, owing to effective phosphate linkages between lignin and chitosan that limited chain mobility and water uptake (Ahmadi et al., 2015). Similar reductions in swelling due to phosphate and covalent crosslinking were reported by Wang et al. (2019), where STMP-crosslinked chitosan/methylcellulose films showed ∼139% swelling vs. ∼196% in controls (Wang et al., 2019), and by Encinas-Basurto et al. (2021), where increasing genipin concentration from 1% to 5% reduced swelling from 205% to 41% (Encinas-Basurto et al., 2025). Hydrogels crosslinked with 1.5%(w/v) STMP exhibited higher swelling (31.85 ± 2.32%) on Day 2 which revealed that further increase in STMP concentration did not lower swelling index but rather has compromised the flexibility of the film. Thus, 1% (w/v) STMP was identified as optimal with superior polymeric integrity and reduced equilibrium swelling.

**Fig. 1.**
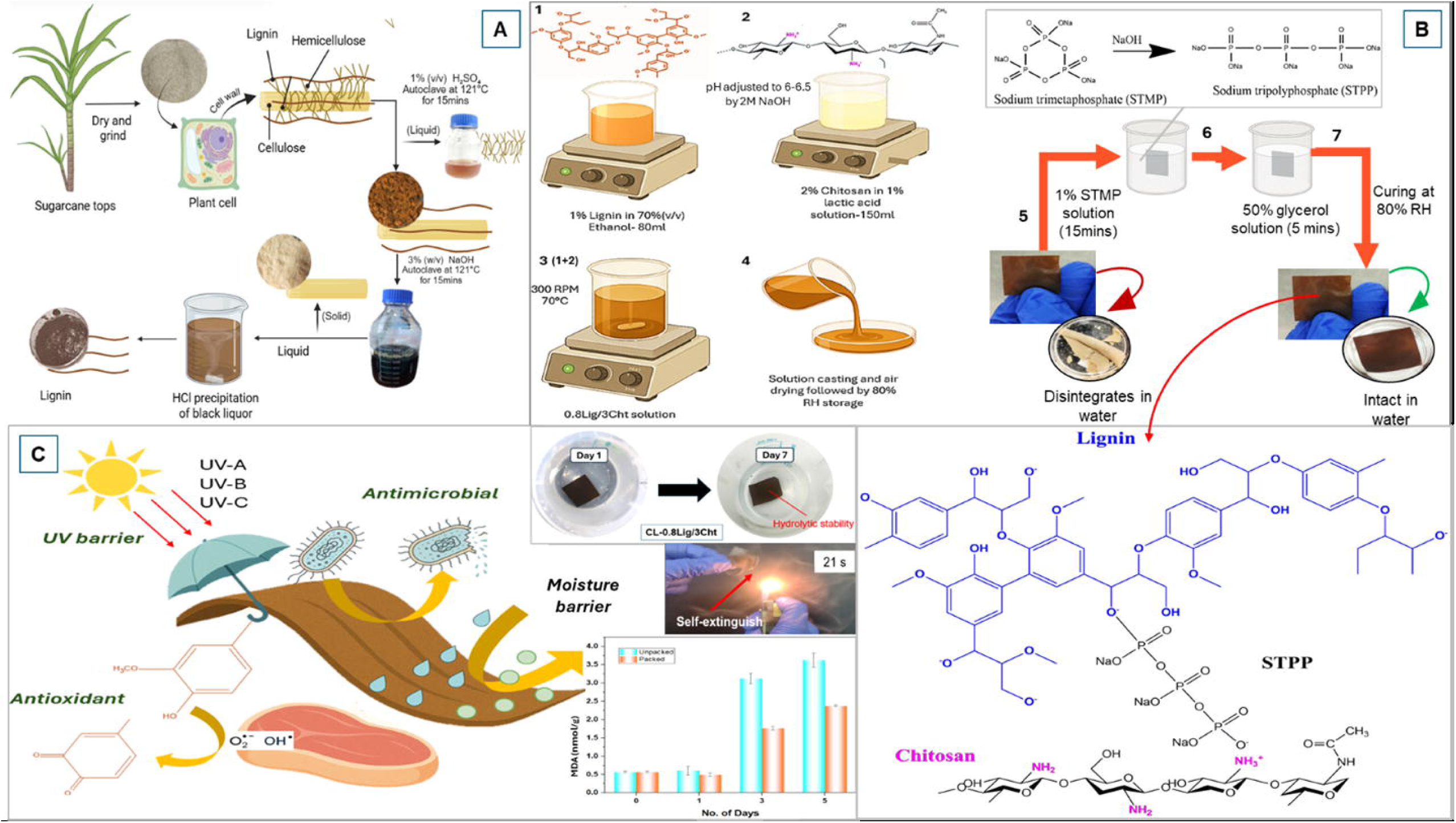
(A) Effect of STMP concentration on hydrogel swelling, **(B)** Morphology of non-crosslinked (softened) vs. crosslinked (intact) films, **(C)** Influence of STMP dipping time on swelling (% w/w), **(D)** WVTR and contact angle of 0.8Lig/3Cht films and CL-0.8Lig/3Cht films, **(E)** Benchmarking of WVTR and swelling performance against reported biopolymer-based packaging films, and **(F)** Percentage swelling of CL-0.8Lig/3Cht films in food simulants on Day 7.

### 3.2 Effect of STMP dipping time on swelling behaviour

To evaluate the effect of crosslinking duration on hydrogel performance, CL-0.8Lig/3Cht films were immersed in 1% (w/v) STMP solution for 15–75 min, and swelling was measured after 1 h and 24 h of water immersion (Fig. 1C). Swelling studies revealed minimal variation across all durations i.e. 15 min (24.14 ± 2.02% at 1 h; 22.69 ± 1.74% at 24 h), 30 min (26.22 ± 0.57% at 1 h; 23.44 ± 0.61% at 24 h), 45 min (26.71 ± 3.42% at 1 h; 23.64 ± 3.27% at 24 h), 60 min (23.40 ± 1.50% at 1 h; 20.92 ± 0.94% at 24 h), and 75 min (26.52 ± 0.83% at 1 h; 23.87 ± 1.34% at 24 h). Two-factor ANOVA confirmed no significant effect of dipping time at either 1 h (p-value = 0.39) or 24 h (p = 0.45). These results indicate that prolonged STMP exposure does not increase crosslinking density, likely due to a limited number of reactive sites or rapid initial reaction kinetics (Lin-Gibson et al., 2003) Therefore, 15 min of STMP dipping is sufficient for achieving optimal and saturated crosslinking, enabling efficient film fabrication without extended processing time.

### 3.3 Water vapor transmission rate (WVTR)

The CL-0.8Lig/3Cht hydrogel film exhibited a remarkable moisture-barrier performance, achieving a WVTR of 24.3 g/m²/day (Fig. 1D) indicating 2.69-folds reduction compared to the non-crosslinked control. This improvement confirms the formation of a dense, moisture-restricting network that minimizes dehydration and microbial growth, both of which is critical for preserving meat freshness and shelf life. Typical biopolymer films show high WVTR due to their hydrophilic and porous nature. For example, nanocellulose-based films exhibit WVTRs from within a range of 696 to 9600 g/m²/day (H. Tayeb et al., 2020) (Bedane et al., 2015), while chitosan composites range from 120.7-167.8 g/m²/day (Wahba et al., 2025). Chemically modified starch systems can lower WVTR from 1453 to 48 g/m²/day (La Fuente Arias et al., 2023), and boronated chitosan achieves as low as 3–5 g/m²/day (Mylkie et al., 2025). However, these studies lack swelling and solvent-resistance evaluations, limiting direct comparison with the present work. Collectively, this evidence establishes the CL-0.8Lig/3Cht film as a superior moisture-barrier film, highlighting its strong potential for sustainable food packaging. The CL-0.8Lig/3Cht film exhibits a highly dense and compact microstructure which directly contributes to its lower swelling ratio and reduced water vapor transmission rate (WVTR) compared to reported biopolymeric films (Fig. 1E).

### 3.4 Wettability

Static sessile drop contact angle analysis was performed to assess the wettability of 0.8Lig/3Cht and CL-0.8Lig/3Cht films (Fig. 1D). The 0.8Lig/3Cht film showed contact angles of 90.5° (left) and 91.2° (right). This value is consistent with reported chitosan–lignin films (∼85–95°) where lignin’s aromatic domains enhance hydrophobicity (Crouvisier-Urion et al., 2016). In contrast, the CL-0.8Lig/3Cht film exhibited lower angles of 69.7° (left) and 73.6° (right) indicating increased surface hydrophilicity after STMP crosslinking. Similar trends have been observed for citric-acid- and vanillin-crosslinked chitosan films, where newly introduced polar groups orient toward the surface (Rani et al., 2025). Despite higher wettability, the CL-0.8Lig/3Cht films retained structural integrity during prolonged aqueous immersion, unlike typical hydrophilic biopolymers that highly swell or dissolve. The decoupling of surface hydrophilicity and bulk water resistance marks a key advancement in interfacial design. Crosslinking reorganizes the polymer network, orienting polar groups at the surface while maintaining a dense core that resists swelling and dissolution. This balance of surface adhesion and structural stability makes CL-0.8Lig/3Cht film a strong candidate for high-moisture food (Lin et al., 1996a) packaging applications.

### 3.5 Stability in food simulants

Fig. 1F depicts the interaction of the CL-0.8Lig/3Cht films with various food simulants, showing leaching values of 5.07 ± 1.18% in 3% (v/v) acetic acid, 4.57 ± 0.13% in water, and 4.54 ± 1.03% in 10% (v/v) ethanol. The marginal variation among these values demonstrates that the films maintained strong structural stability and solvent resistance, with their densely crosslinked matrix effectively restricting lysis and leaching. This stability surpasses conventional biopolymer films, which often degrade under similar conditions. Pei et al. (2025) reported that chitosan-based composite films suffered structural collapse and essential oil migration in 3% (v/v) acetic acid (Lin et al., 1996b), while others have shown that chitosan films lose integrity under high humidity (∼98% RH) due to swelling and gel formation(Orcutt et al., 2025). In contrast, the CL-0.8Lig/3Cht films-maintained leaching below 5% across all simulants, including acidic media, confirming a highly efficient crosslinking mechanism that prevents acid-catalyzed dissolution of biopolymers. Furthermore, unlike conventional chitosan film fabrication requiring post-processing neutralization with NaOH (Llanos et al., 2015), CL-0.8Lig/3Cht films achieved stability in food simulants through a single crosslinking process offering enhanced durability, scalability, and environmental compatibility for sustainable food-contact applications.

### 3.6 Key interactions in hydrogel films

XPS analysis of 0.8Lig/3Cht and CL-0.8Lig/3Cht films is shown is Fig. 2. In the C 1s spectra of Fig. 2 (A and B), three major components were observed where a dominant peak at 284.6 eV corresponding to C–C/C–H bonds from lignin’s aromatic rings and chitosan’s glucosamine backbone. Another peak at 286.1 eV is assigned to C–N/C–O bonds from hydroxyl and amine groups and a higher binding energy peak at 288.1 eV is attributed to C=O groups from acetylated chitosan residues and lignin carbonyls.

**Fig. 2.**
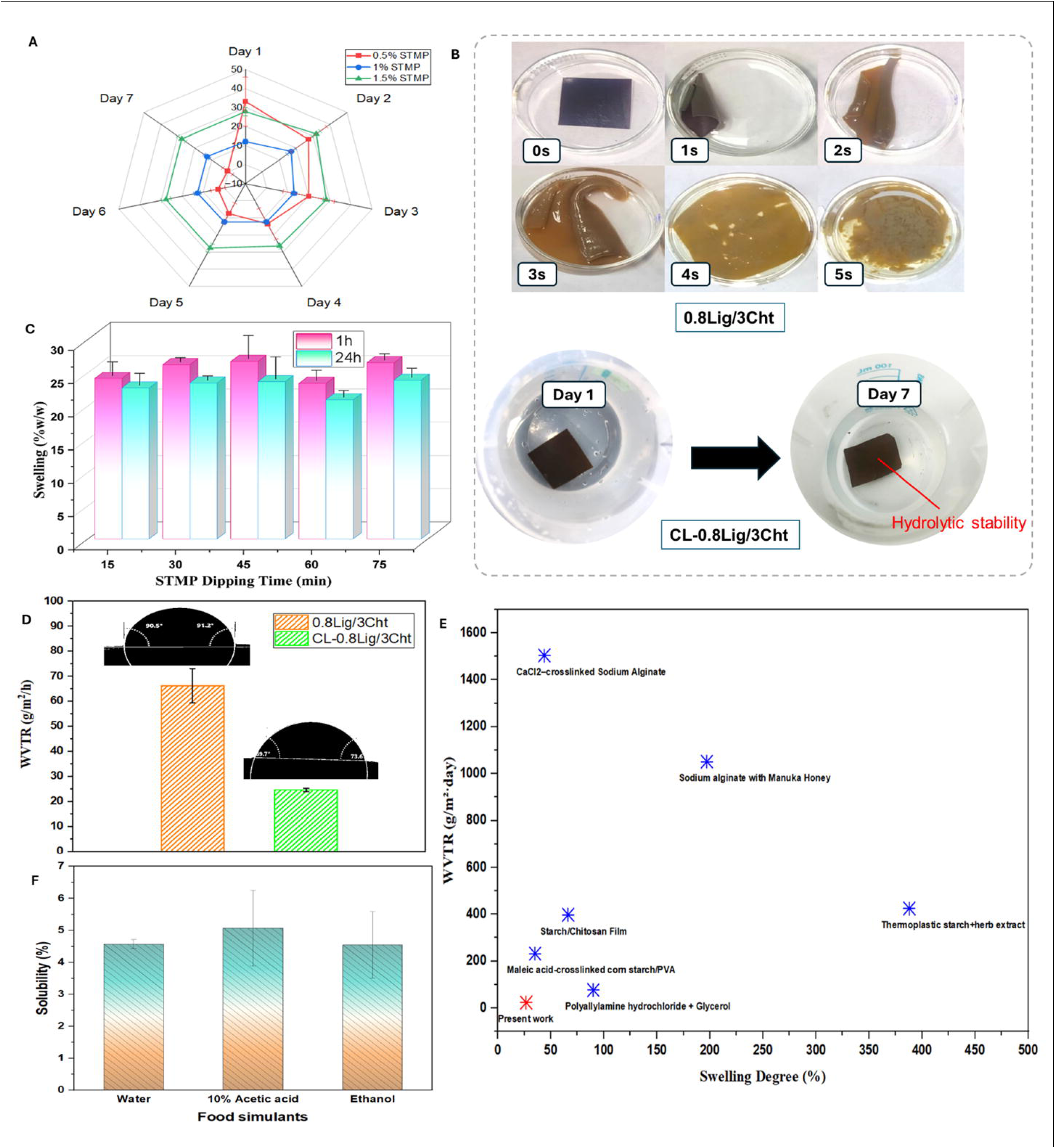
XPS analysis of 0.8Lig/3Cht and CL-0.8Lig/3Cht films **(A, B)** C 1s spectrum depicting different carbon bonding states, **(C, D)** N 1s spectrum showing nitrogen functionalities before and after crosslinking, and **(E, F)** P 2p spectrum verifying phosphorus integration into the film matrix.

In the N 1s spectra in Fig. 2 (C and) peaks at 398.8 eV and 400.8 eV correspond to non-protonated amines (–NH[) from deacetylated chitosan (Yue et al., 2018) and amide groups (–NH–C=O) from N-acetylglucosamine units (Pradal et al., 2011), respectively. The reduction in the 400.8 eV amide peak intensity in the crosslinked film suggests partial modification of acetylated sites during phosphate incorporation.

The P 2p region displayed a prominent peak at 130.1 eV (Fig. 2 F), indicating the presence of low-valent phosphorus species formed during STMP crosslinking (Pradal et al., 2011). This shift reflects partial reduction of P (+5) species, with phosphorus forming P–O–N ionic interactions with chitosan amines(Pati et al., 2011) and P–O–aryl covalent linkages with lignin’s phenolic hydroxyls, producing stable phosphate esters (Lignin–O–P) (Chaudhari et al., 2024)(Fig. 3). Although such electrostatic P–O–N interactions are typically unstable in aqueous environments and prone to degradation, the remarkable water stability observed here indicates the formation of covalent, hydrolytically stable phosphate ester bonds. These interactions collectively enhance the structural integrity of the film with exceptional water resistance.

**Fig. 3.**
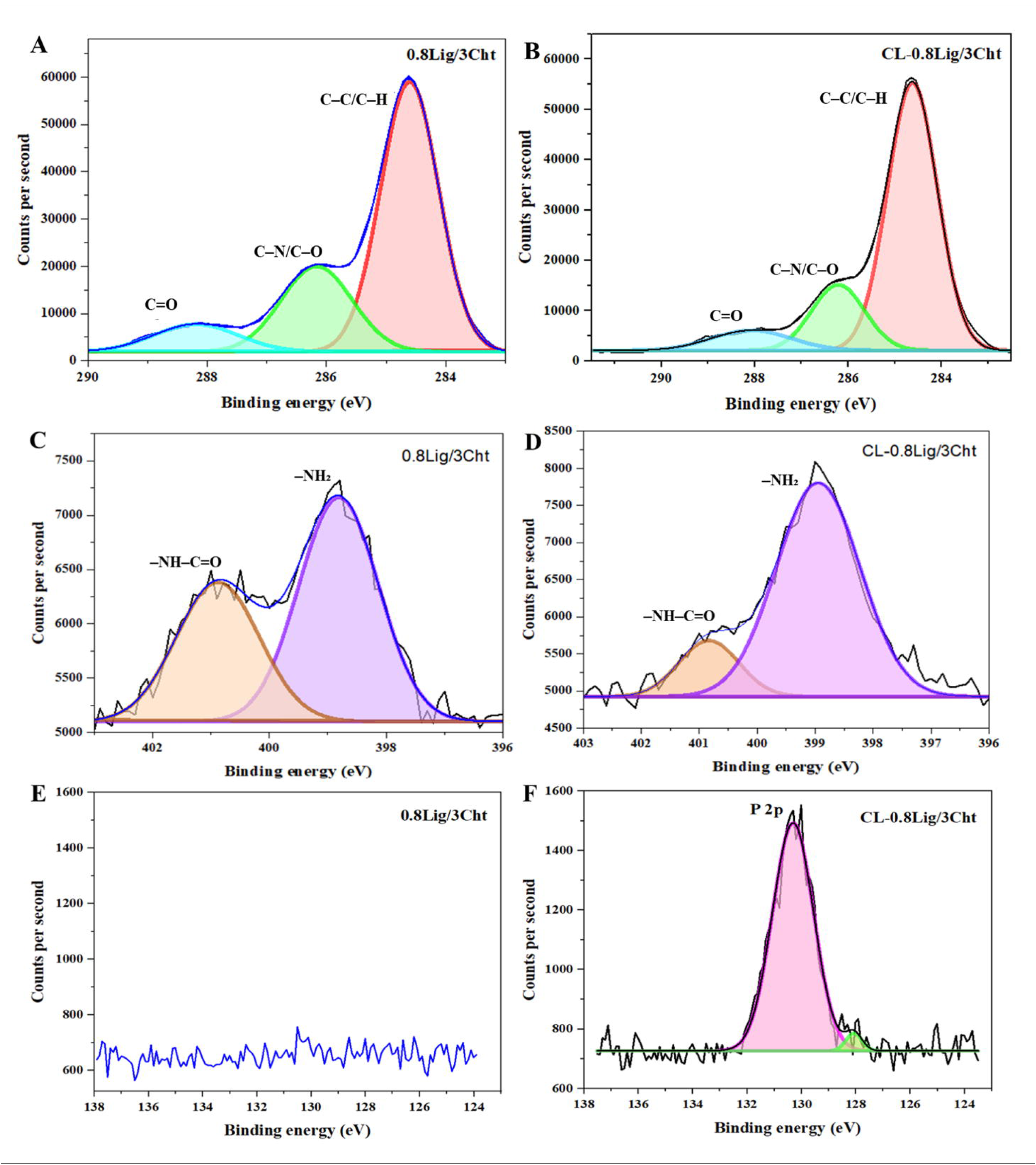
Proposed interaction of the STMP-crosslinked lignin–chitosan hydrogel network illustrating phosphate amino and ester bond between chitosan and lignin phenolic sites.

### 3.7 UV barrier activity

UV protection is vital in meat packaging, as UV exposure accelerates lipid oxidation, protein denaturation, and myoglobin degradation, causing rancidity, discoloration, and nutrient loss. UV transmittance analysis (200–400 nm) showed that the CL-0.8Lig/3Cht film maintained strong UV-B shielding, with a blocking efficiency of 98.38%, confirming that crosslinking did not compromise its UV-protective function (Fig. 4A). The SPF values of 62.60 in 0.8Lig/3Cht and 61.59 in CL-0.8Lig/3Cht (Fig. 4B) further indicated negligible loss in UV resistance after STMP modification. These results align with lignin-reinforced systems, where cellulose/lignin composites blocked 82.09% UV-B and 61.93% UV-A with 5 wt % of lignin (Li et al., 2024). The pure chitosan film showed 23.39-fold lower UV-B blocking, emphasizing lignin’s aromatic and phenolic π–π* and n–π* transitions as the main contributors to UV attenuation (Goliszek-Chabros et al., 2025). The absorption peaks at ∼310 nm (Cα═Cβ linkages), 280–240 nm (hydroxyl groups), and 200–210 nm (aromatic rings) shown in Table S1 confirm these transitions. Unlike conventional LDPE films requiring benzotriazole stabilizers that risk leaching (Khare et al., 2025), lignin in the CL-0.8Lig/3Cht matrix remains immobilized, providing a non-toxic, food-safe, and durable UV-shielding packaging alternative.

**Fig. 4.**
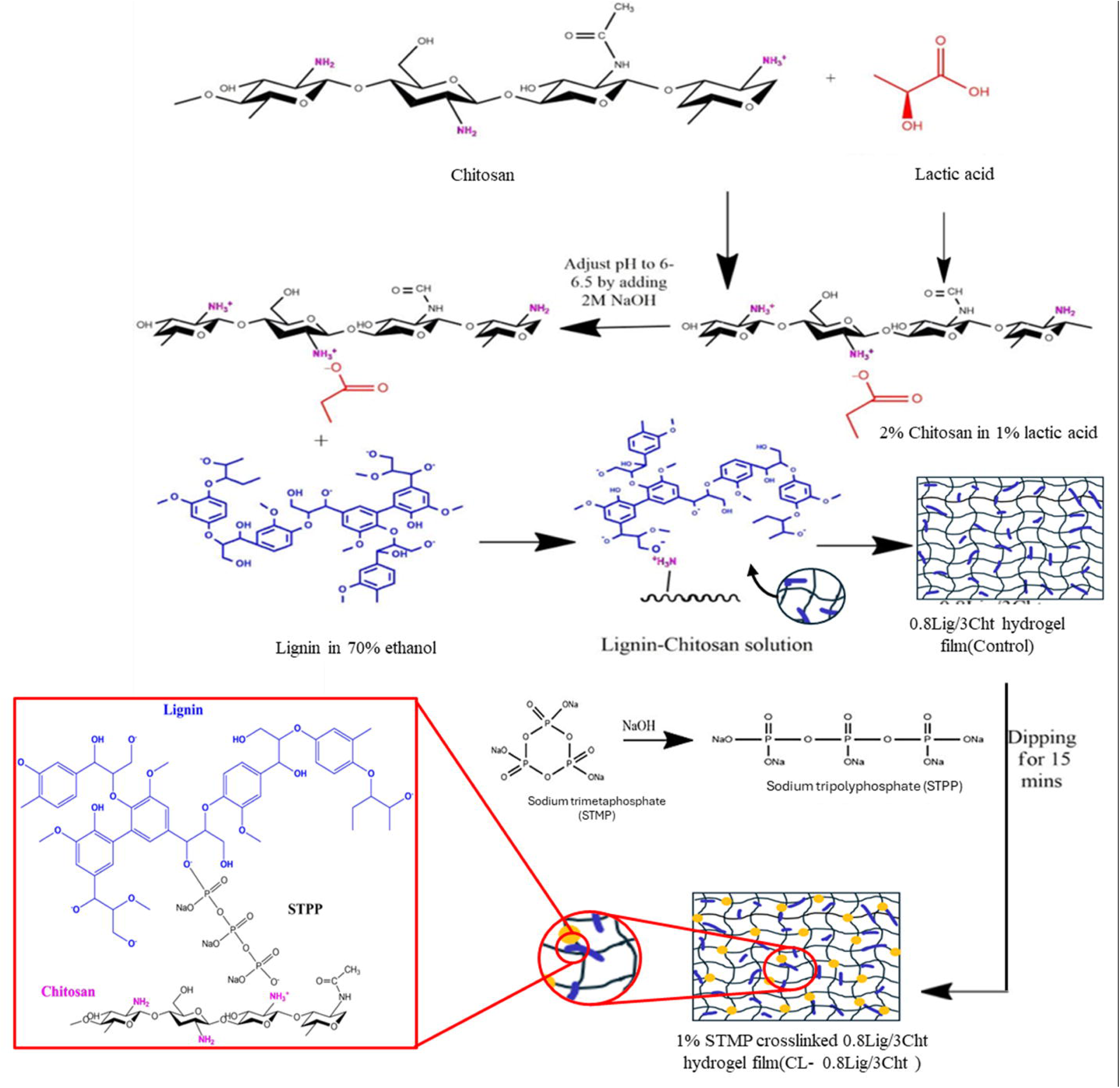
(A) UV–visible light transmittance spectra of STMP-crosslinked (CL-0.8Lig/3Cht) and non-crosslinked (0.8Lig/3Cht) hydrogel films, **(B)** Comparison of Sun Protection Factor (SPF) and UV-B blocking percentage among 3Cht, 0.8Lig/3Cht, and CL-0.8Lig/3Cht films, **(C)** Energy level transitions and associated lignin structural features, and **(D)** Antimicrobial activity of the CL-0.8Lig/3Cht films against DH5α.

### 3.8 Antimicrobial Activity

The antimicrobial performance of the CL-0.8Lig/3Cht hydrogel film was assessed against *E. coli* DH5α using the agar diffusion assay shown in Fig. 4D. A clear inhibition zone of 7.0 ± 0.5 mm, slightly exceeding the film disc diameter of 6.0 mm, confirmed its antibacterial efficacy. This activity, though moderate, highlights the film’s intrinsic antimicrobial potential without reliance on synthetic additives. The observed effect is attributed to the combined action of chitosan’s cationic amine groups that disrupt bacterial membranes, and lignin’s phenolic hydroxyls that can induce oxidative stress(Garg and Avanthi, 2025). The STMP crosslinking did not mask these reactive groups. Instead, it preserved their accessibility within the matrix, enabling sustained bioactivity while enhancing the film’s structural integrity. The retention of antimicrobial function post-crosslinking supports the film’s application in food packaging via protection against food spoilage causing microbes.

### 3.9 Radical scavenging activity

Radical scavenging activity (RSA) is essential in active food packaging to prevent oxidative degradation that leads to quality loss in perishable foods such as meats and high-fat products (Romani et al., 2020). The CL-0.8Lig/3Cht hydrogel films exhibited a higher RSA of 89.89%, representing 1.2-folds increase relative to 0.8Lig/3Cht films. This strong antioxidant effect arises from lignin’s phenolic hydroxyl and methoxy groups, which stabilize DPPH radicals through resonance-stabilized phenoxy intermediates. Additionally, STMP-induced phosphate crosslinking enhanced RSA via free radical quenching mechanism consistent with previous findings on phosphorylated polysaccharides(Xie et al., 2020). Similar studies reported 72.5% RSA for phosphorylated polysaccharides (Cui et al., 2025)and an increase from 37.8% to 65.9% after phosphorylation (Zhang et al., 2022). The synergistic interaction between lignin’s phenolic architecture and phosphate functionalities endowed the film with superior antioxidant capacity which is a key attribute for delaying lipid oxidation, pigment degradation, and quality loss during storage.

### 3.10 Mechanical robustness of hydrogel films

The mechanical strength of the hydrogel films remarkably improved after STMP-induced crosslinking. The CL-0.8Lig/3Cht film exhibited a tensile strength of 2364.79 MPa i.e.,41-folds increase related to 0.8Lig/3Cht film (Fig. 5A), indicating the formation of a dense phosphate-linked 3D network that restricts polymer chain slippage and enhances stress transfer. Crosslinking-induced reinforcement is well-documented in biopolymer systems, though typically of lower magnitude. Qin et al. (2022) reported a strength increase from 42.6 MPa to 80.7 MPa with elongation decreasing from 23.7 % to 5.6 % in tannic acid–modified chitosan films (Qin et al., 2022). Wang et al. (2019) achieved 294 MPa in PLA–cellulose nanocrystal hybrid films (Fitriani et al., 2023). Glycerol, used as a plasticizer improved flexibility by increasing molecular mobility while slightly reducing tensile strength, consistent with starch–catechin–nanocrystal films. In starch–catechin–starch nanocrystal films, glycerol incorporation increased elongation at break while slightly reducing tensile strength due to improved molecular mobility and relaxation within the polymer network (Sessini et al., 2016). The present formulation of CL-0.8Lig/3Cht film was specifically designed to overcome the conventional trade-off between strength and flexibility through inclusion of appropriate STMP and glycerol proportions in the total polymer matrix to maintain 31.49 % elongation alongside its high strength. Since materials with elongation < 5 % are considered brittle (Astakhov, 2018), the CL-0.8Lig/3Cht film demonstrates balanced flexibility and robustness suitable for stress-bearing food packaging applications where both durability and functional adaptability are essential.

**Fig. 5.**
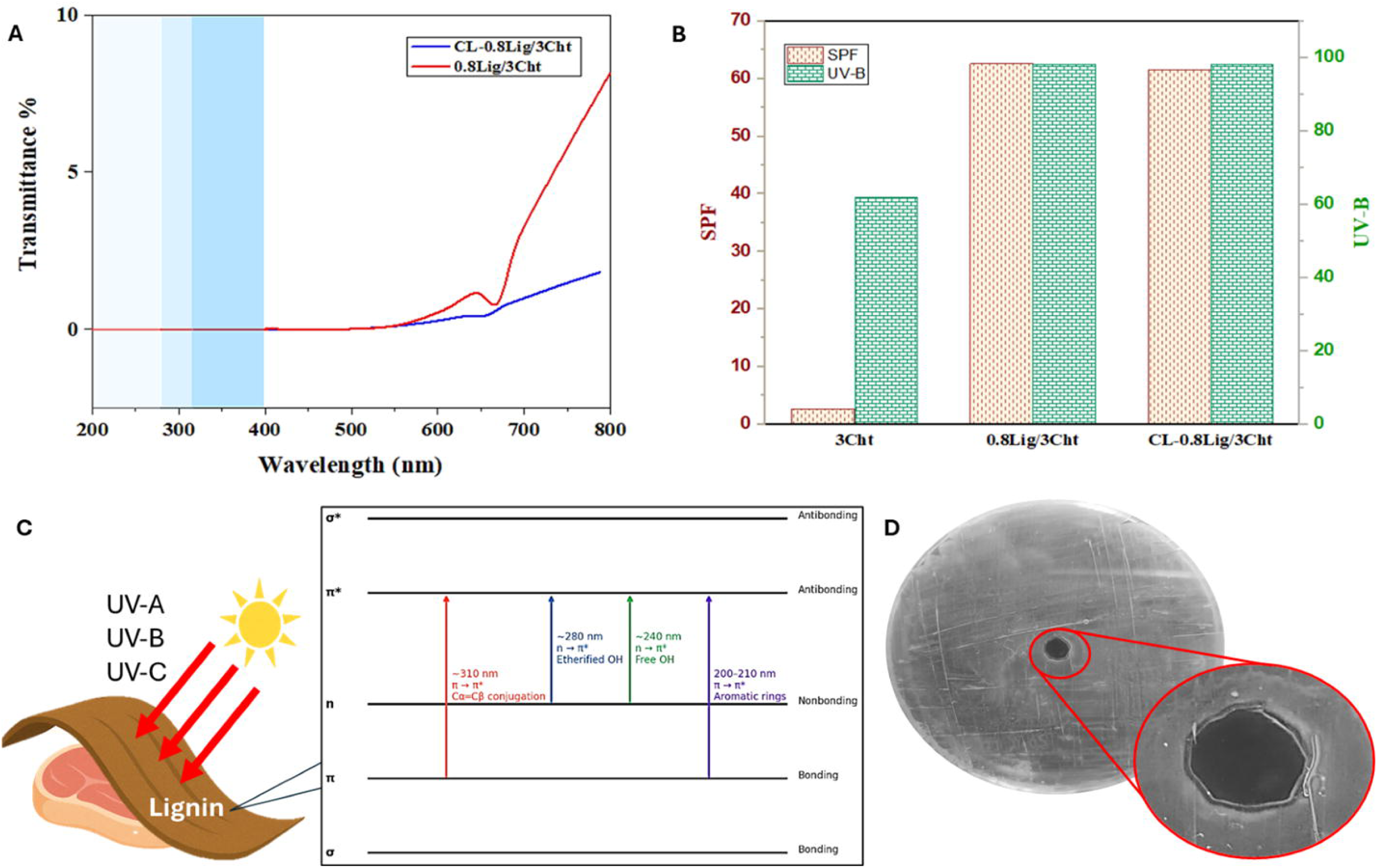
(A) Tensile stress–strain profiles of 0.8Lig/3Cht and CL-0.8Lig/3Cht films. (B) Visual demonstration of the mechanical durability of the CL-0.8Lig/3Cht film showing its flexibility (left) and ability to withstand a 250 g load without tearing (right), (C) Anti-freezing, and (D) Anti-wrinkle performance of the CL-0.8Lig/3Cht film.

### 3.12 Anti-freezing and anti-wrinkle property of hydrogel films

The anti-freezing property of packaging films is crucial for maintaining meat quality during cold storage and transport. Conventional bio-based materials often become brittle or lose barrier integrity under freezing conditions leading to moisture loss and spoilage. In contrast, the CL-0.8Lig/3Cht hydrogel films (Fig. 5C) exhibited flexibility under low-temperature conditions, reflecting strong anti-freezing capability. This property arises from multiple synergistic mechanisms of alkali lignin, chitosan, and glycerol. Glycerol within the crosslinked matrix acts as a cryoprotectant by forming hydrogen bonds with water, thereby inhibiting ice crystallization and lowering the freezing point (Alba-Simionesco et al., 2022). Alkali lignin enhances hydrogen bonding with glycerol and increases network density, which minimizes glycerol migration and promotes water retention (Han et al., 2023). Moreover, chitosan is reported to show stability to freeze–thaw by limiting water mobility and delaying ice recrystallization through intermolecular hydrogen-bonding interactions (Tang et al., 2025). These effects impart superior flexibility and structural resilience to the hydrogel film under sub-zero environments, underscoring its potential for storage and transportation of frozen meat.

Another crucial attribute in flexible packaging materials is their resistance to permanent deformation under mechanical stress. A film that can sustain bending, folding, or compression without developing irreversible creases or cracks ensures not just physical protection of food items but also long-term material integrity and consumer appeal. Fig. 5D highlights the film’s anti-wrinkle performance, as it can be observed that after being crumpled and left undisturbed, the films on opening regained their initial shape without permanent distortions. The inherent elasticity of the films prevented structural damage, allowing them to maintain their mechanical integrity. Since no visible surface cracks were observed, it can be substantiated that the films exhibit enhanced durability, which is crucial for protecting food products from physical damage and ensuring prolonged shelf life.

### 3.13 Thermogravimetric analysis (TGA)

Thermogravimetric analysis (TGA) was conducted to assess the thermal stability of lignin– chitosan hydrogel films and the influence of STMP crosslinking on thermal degradation resistance (Fig. 6). The initial endothermic peak between 70–100 °C corresponds to the evaporation of entrapped water, with the evaporation temperature shifting from 70 °C in the chitosan film (control) to 100 °C in 0.8Lig/3Cht and CL-0.8Lig/3Cht films, indicating restricted water mobility due to the denser crosslinked matrix. A second endothermic transition was observed at around 160 °C in chitosan film, 80 °C in 0.8Lig/3Cht and 130 °C in CL-0.8Lig/3Cht. The decrease in melting temperature with lignin addition reflects its plasticizing effect which disrupts chitosan’s crystalline packing and forms lower-melting defective crystallites (Rim and Runt, 1984). Conversely, STMP crosslinking raised the melting point to 130 °C which is consistent with prior findings for citric acid–crosslinked chitosan film (Guerrero et al., 2019). The enhanced thermal stability of CL-0.8Lig/3Cht film indicates resilience to temperature fluctuations during processing, transportation, and storage, supporting its suitability for active packaging applications (Pérez-Córdoba et al., 2018).

**Fig. 6.**
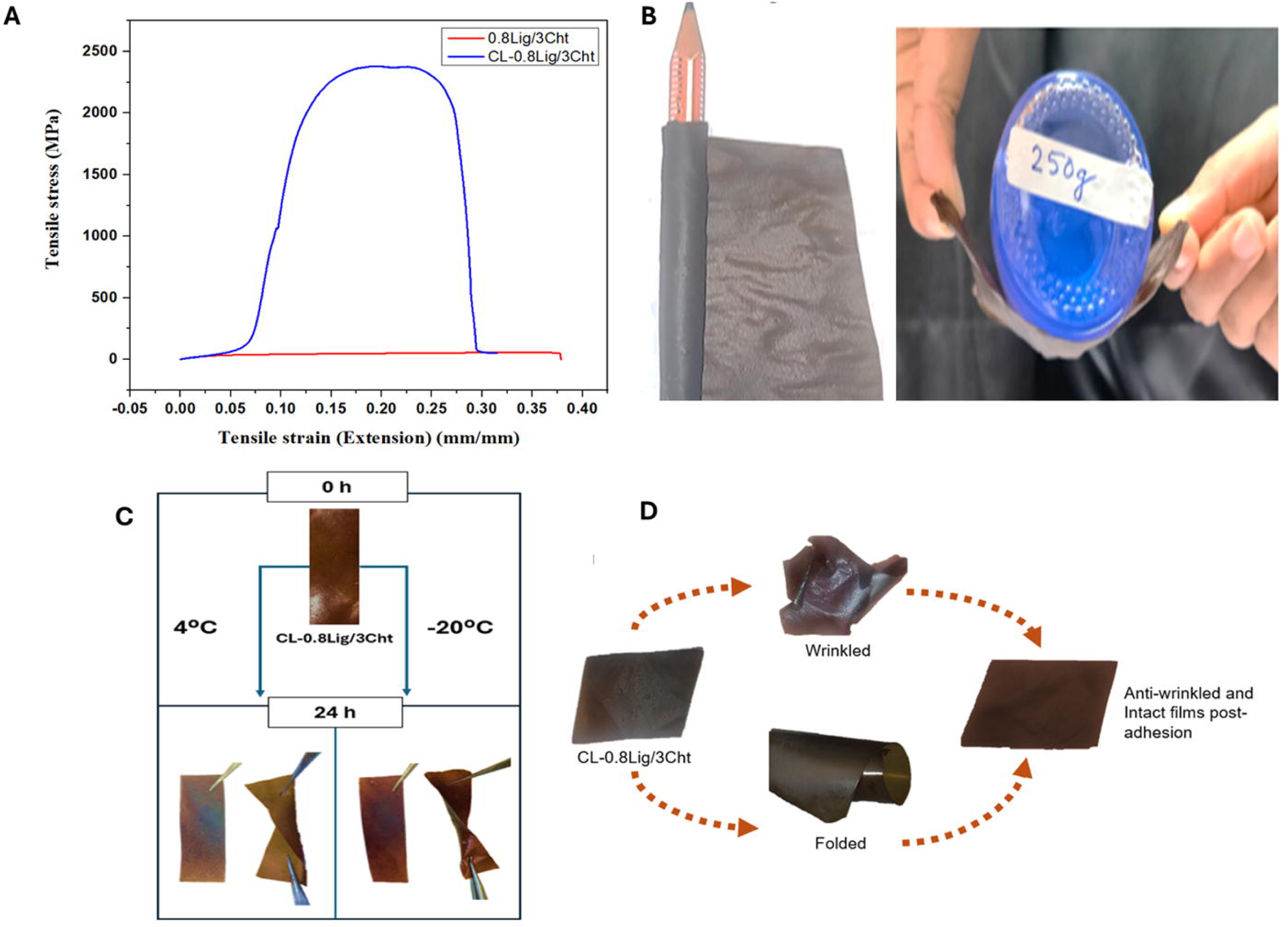
Thermogravimetric analysis (TGA) curves of 3Cht, 0.8Lig/3Cht, and CL-0.8Lig/3Cht hydrogel films illustrating their thermal degradation profiles.

### 3.14 Differential scanning calorimetry (DSC) analysis

DSC analysis of chitosan, 0.8Lig/3Cht and CL-0.8Lig/3Cht films is shown in Fig. 7 (A to C) which is crucial for active meat packaging films as it identifies thermal transitions influencing film stability, integrity, and performance during sealing and storage. The DSC thermogram of pure chitosan film displayed a broad endothermic peak near 100 °C, corresponding to the evaporation of bound and free water from its hydrophilic matrix (Janik et al., 2021), followed by a major degradation event at 196 °C attributed to the decomposition of amine units. Incorporation of lignin and phosphate crosslinking (via STMP) significantly altered the thermal behaviour of 0.8Lig/3Cht and CL-0.8Lig/3Cht films. Both exhibited an endothermic peak between 70–100 °C due to entrapped water evaporation. The semicrystalline chitosan imparted flexibility, while amorphous lignin enhanced thermal rigidity through aromatic π-conjugation within its phenylpropanoid framework by elevating their onset decomposition temperatures (Belouadah et al., 2024). Lignin likely acts synergistically with the hydrogel matrix to disrupt crystallinity while enhancing thermal resistance (Li et al., 2023). Crosslinking with STMP shifted the primary endothermic peak from ∼112 °C to ∼135 °C, indicating improved thermal stability through a denser, moisture-restrictive network. The secondary transition, linked to polymer softening, also shifted from 80 °C in the 0.8Lig/3Cht to 130 °C in CL-0.8Lig/3Cht film, confirming enhanced rigidity and structural integrity. These findings are consistent with char residue and LOI data (Table 1) which demonstrate that STMP crosslinking improves both thermal and flame-retardant properties of the hydrogel matrix.

**Fig. 7.**
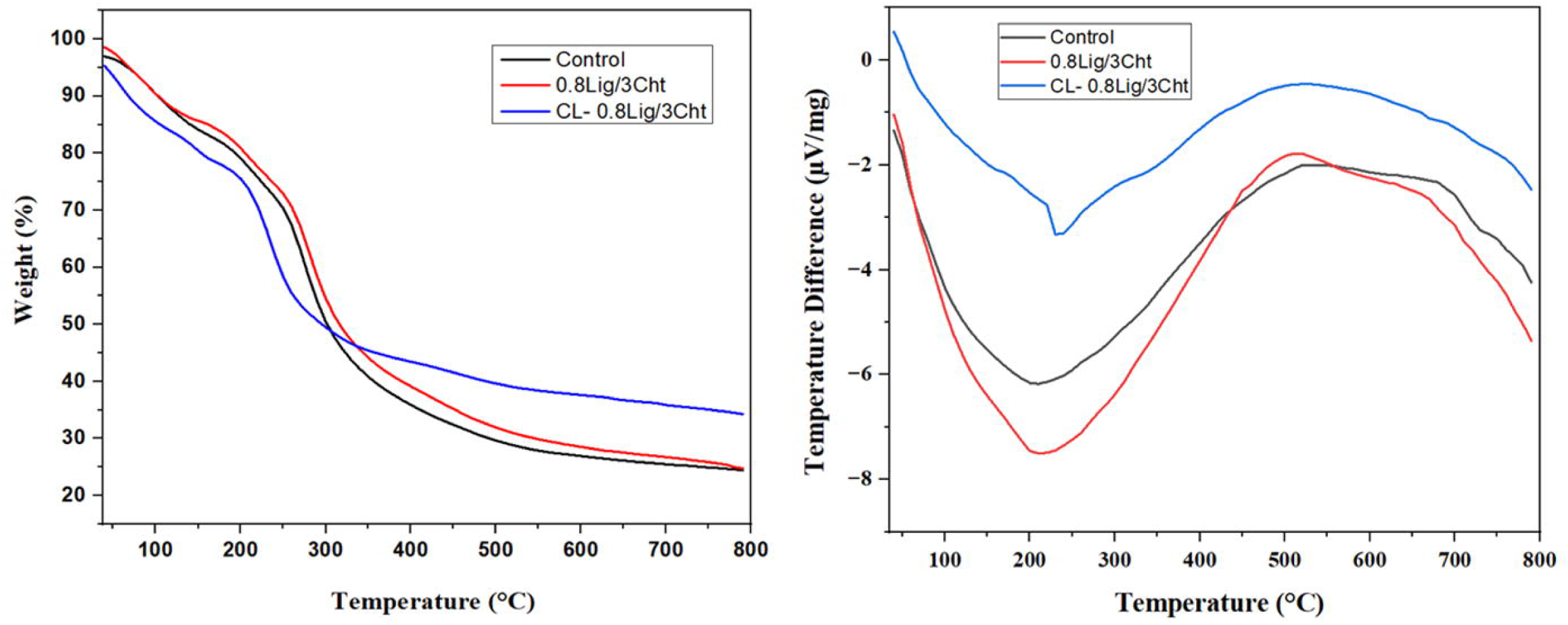
Differential scanning calorimetry (DSC) thermograms of **(A)** 3Cht, **(B)** 0.8Lig/3Cht, and **(C)** CL-0.8Lig/3Cht films, and **(E, F)** Flammability test depicting delayed ignition and rapid self-extinguishing behaviour of CL-0.8Lig/3Cht films compared to 3Cht.

**Table 1.** Char residue, limiting oxygen index (LOI), and corresponding flame resistance level of the hydrogel film samples.

| Sample | Char Residue % | Calculated LOI (%) | Flame Resistance Level |
| --- | --- | --- | --- |
| 3Cht | 24.47 | 7.29 | Moderate / Self-extinguishing |
| 0.8Lig/3Cht | 24.76 | 27.4 | Moderate / Self-extinguishing |
| CL-0.8Lig/3Cht | 34.28 | 31.21 | High / Strong flame resistance |

#### 3.15.1 Surface architecture and elemental analysis of hydrogel films

To elucidate the surface architecture and crosslinking-induced structural development of the hydrogel films, SEM micrographs were obtained across magnifications of 50 µm, 5 µm, and 1 µm for 0.8Lig/3Cht film, and CL-0.8Lig/3Cht film with and without glycerol (Fig. S1). Upon lignin incorporation (0.8Lig/3Cht), pronounced surface roughness and microstructural discontinuities emerged within the chitosan similar to reported studies (Kang et al., 2024)(Rosova et al., 2021). The CL-0.8Lig/3Cht films displayed a denser and more cohesive surface morphology, characterized by homogeneously distributed domains and the absence of phase-separated lignin aggregates, indicating effective molecular-level integration. Further, when glycerol was incorporated into the crosslinked matrix, the film surface exhibited enhanced compactness and smoothness due to its small molecular size that enabled it to penetrate the polymer network (Eslami et al., 2023). Glycerol promotes plasticizing effect which facilitates polymer chain mobility. In contrast, the crosslinked film without glycerol, although still structurally integrated, showed relatively higher micro-roughness, suggesting tighter intermolecular packing in the absence of a plasticizer. This comparative analysis underscores glycerol’s role in modulating the film’s topography while maintaining crosslinking efficiency and mechanical cohesion.

The elemental composition of the films was analysed using EDS coupled with SEM (Fig. 8). All samples exhibited characteristic peaks of carbon (C), oxygen (O), and nitrogen (N), in the chitosan–lignin matrix. Phosphorus (P) is an indicator of STMP crosslinking that appeared only in the crosslinked films (with/without glycerol) thus validating phosphate incorporation. The 0.8Lig/3Cht control (NC) showed minor P content (3.64 wt.%), likely from trace residues, whereas the CL-0.8Lig/3Cht film displayed a markedly higher P level (15.93 wt.%). The CL-G film showed slightly lower P (11.54 wt.%) but higher O content (43.47 wt.% vs. 41.03 wt.% without glycerol) which can be attributed to hydroxyl-rich structure of glycerol. Nitrogen content remained stable across samples, reflecting uniform chitosan presence. These findings, in conjunction with SEM micrographs, reveal successful phosphate-based crosslinking and glycerol incorporation.

**Fig. 8.**
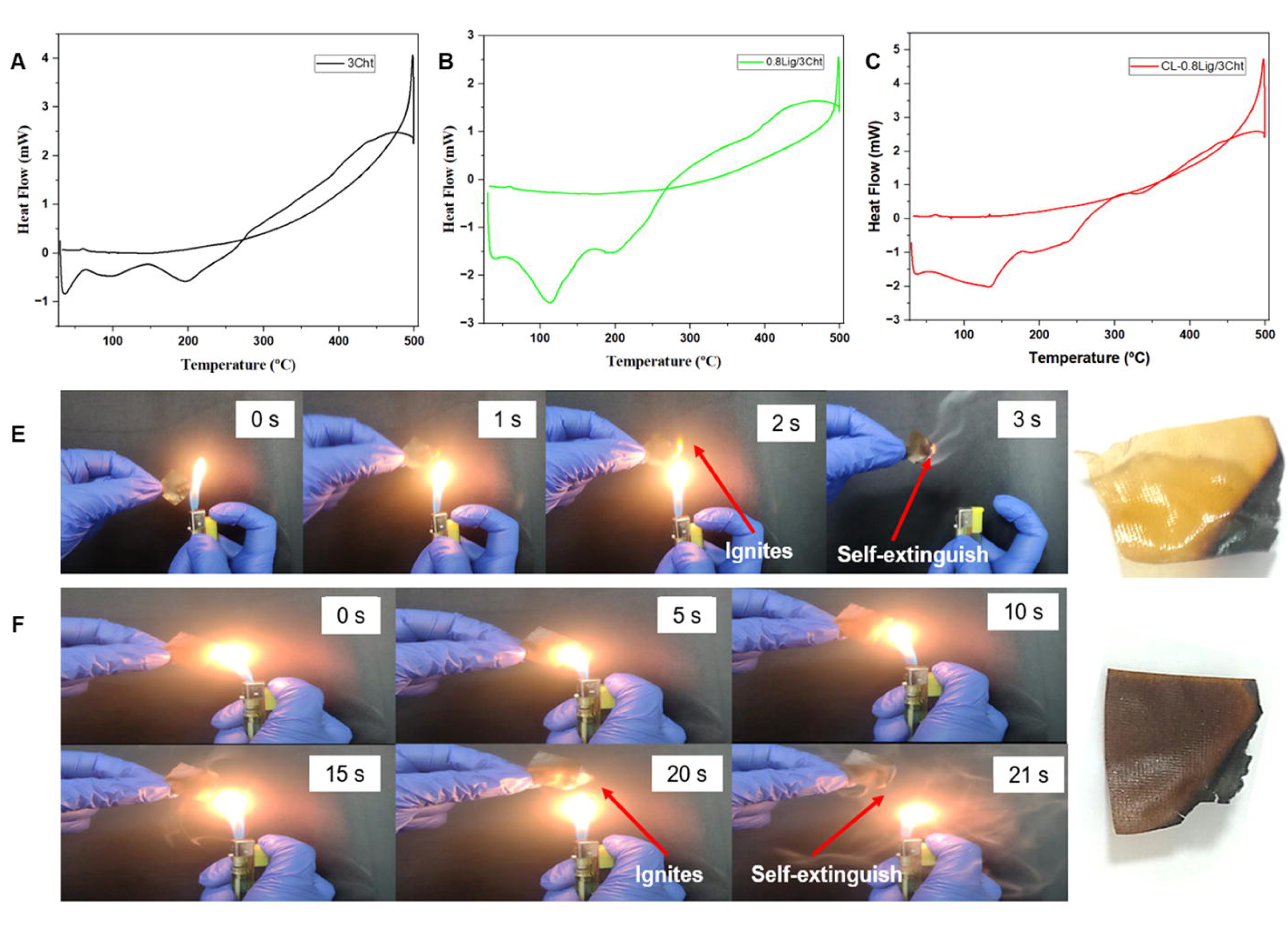
Elemental analysis of NC: Non-crosslinked 0.8Lig/3Cht film, CL: Crosslinked film without glycerol, and CL-G: Crosslinked film with glycerol.

### 3.16 Shelf-life analysis of packaged chicken breast

The chicken breast sample used for the study consisted of 4.63 ±0.7 % (w/w) lipid, 17.12 ± 0.94 % (w/w) protein, 77.96 ± 0.459 % (w/w) moisture, and 2.63 ± 0.7 % (w/w) ash content.

#### 3.16.1 pH variation

The pH variation of packed and unpacked chicken breast samples over five days revealed distinct trends in acidity and spoilage progression (Fig. 9B). Initially, on Day 0, both samples exhibited a high pH of 7.2 which is consistent with the pH of muscle tissue. By Day 1, a sharp decline was observed in both control and test sample, with pH dropping to 5.66 and 5.68 in packed and unpacked conditions. This decline is primarily attributed to post-mortem glycolysis, where glycogen is converted into lactic acid, lowering the pH due to increased acidity. Typically, when meat is preserved under optimum conditions, it should remain fresh with a corresponding pH ranging between 5.0 and 6.5 according to FSSAI. This pH range maintains the texture and microbial stability during storage period. On Day 3, the pH in both samples increased slightly to 6.13 and 6.15 in packed and unpacked samples. This rise is indicative of the onset of microbial activity, where spoilage bacteria start metabolizing proteins, producing basic nitrogenous compounds such as ammonia and amines, which counteract acidity. By Day 5, a significant divergence between the packed and unpacked samples was observed. The pH of packed sample remained relatively stable at 6.1, suggesting a slower spoilage rate, possibly due to limited oxygen exposure and reduced microbial proliferation. However, in the unpacked sample, pH increased sharply to 6.8, indicating extensive microbial spoilage, where bacteria produce alkaline metabolites, accelerating meat deterioration. These results indicate that packaging effectively slows down microbial spoilage, maintaining the pH within typical range of the fresh meat as compared to the unpacked sample.

**Fig. 9.**
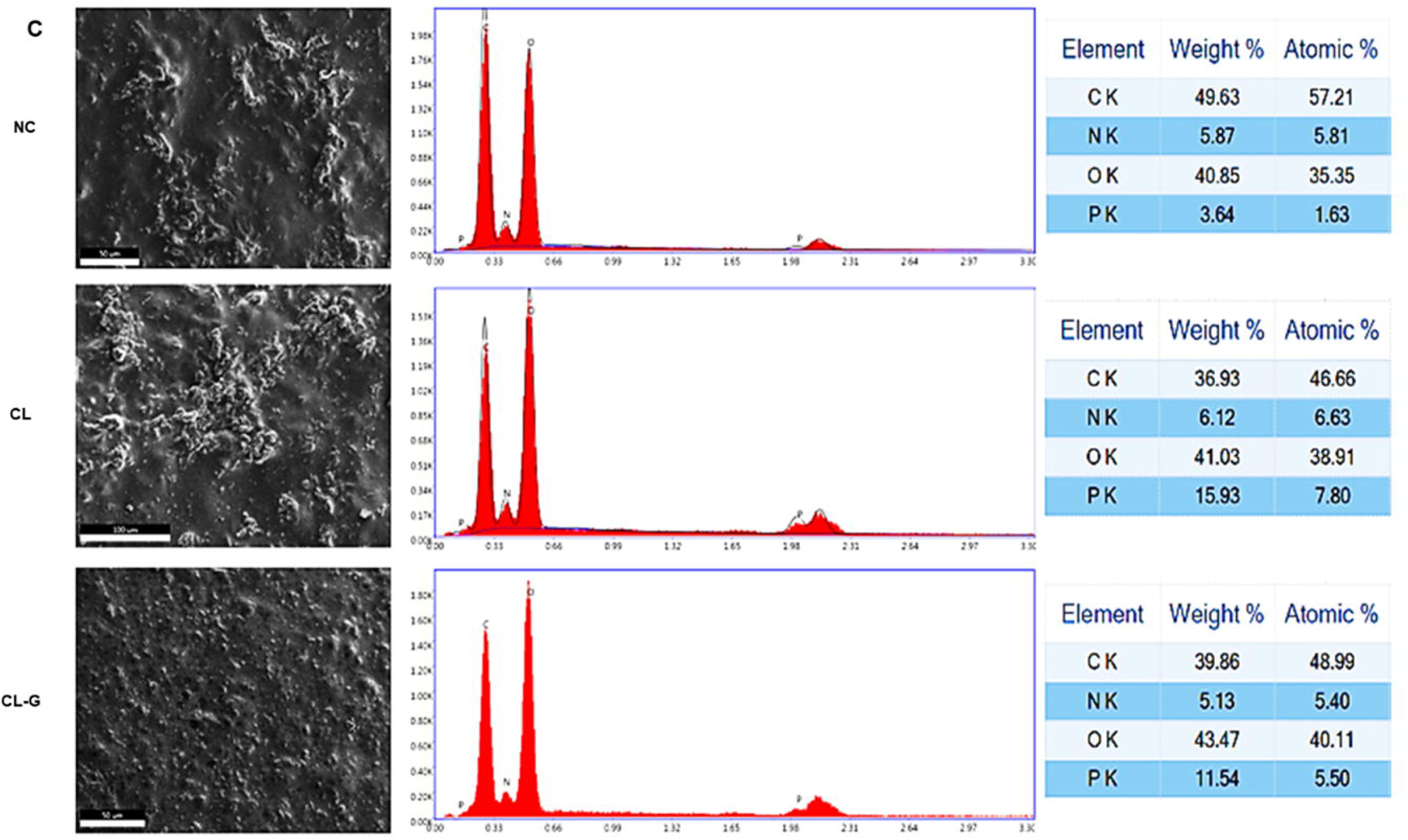
(A) Visual comparison, (B) pH variation, and (C) weight loss (% w/w) of CL-0.8Lig/3Cht packed and unpacked chicken breast over 5 days.

#### 3.16.2 Moisture retention

Moisture loss from the chicken breast surface causes tissue dehydration leading to texture deterioration, accelerated oxidation, and consequent pigment loss (González-López et al., 2023). One of the key functions of meat packaging films is to minimize weight loss by limiting moisture evaporation, thereby preserving overall quality and freshness (Song et al., 2021). Moisture loss was assessed over 5 days to evaluate the CL-0.8Lig/3Cht hydrogel film’s ability to maintain meat juiciness. As shown in Fig. 9C, unpackaged samples exhibited substantially greater moisture loss throughout storage. By Day 5, the unpacked samples lost 75.36 % ± 1.29, whereas the CL-0.8Lig/3Cht-packed samples recorded only 47.60 % ± 0.67. The CL-0.8Lig/3Cht film retained ∼28% more moisture by Day 5 as compared to control sample, thus confirming its efficiency in packaging moist and tender samples which drip water during storage period.

#### 3.16.3 Control on microbial growth

Effective control of microbial proliferation is essential for extending shelf life and ensuring the safety of fresh meat products. Both packed and unpacked groups showed a similar initial microbial load of 6.73 log[[CFU/g on Day 0 (Fig. 10A). By Day 5, the unpacked (control) samples reached 11.95 log[[CFU/g, whereas the CL-0.8Lig/3Cht-packed samples recorded 10.29 log[[CFU/g during storage at 25 °C. A similar trend was observed in a study where poultry stored under fluctuating 0–15 °C cycles exhibited an increase in total viable count from 5.47 to 8.98 log CFU/g after 140 h (∼5.8 days)(Farahnaz Ghollasi-Mood, 2016). Despite the higher storage temperature in this study, the CL-0.8Lig/3Cht film maintained substantial inhibitory activity, suggesting its potential efficacy across a broader temperature range.

**Fig. 10.**
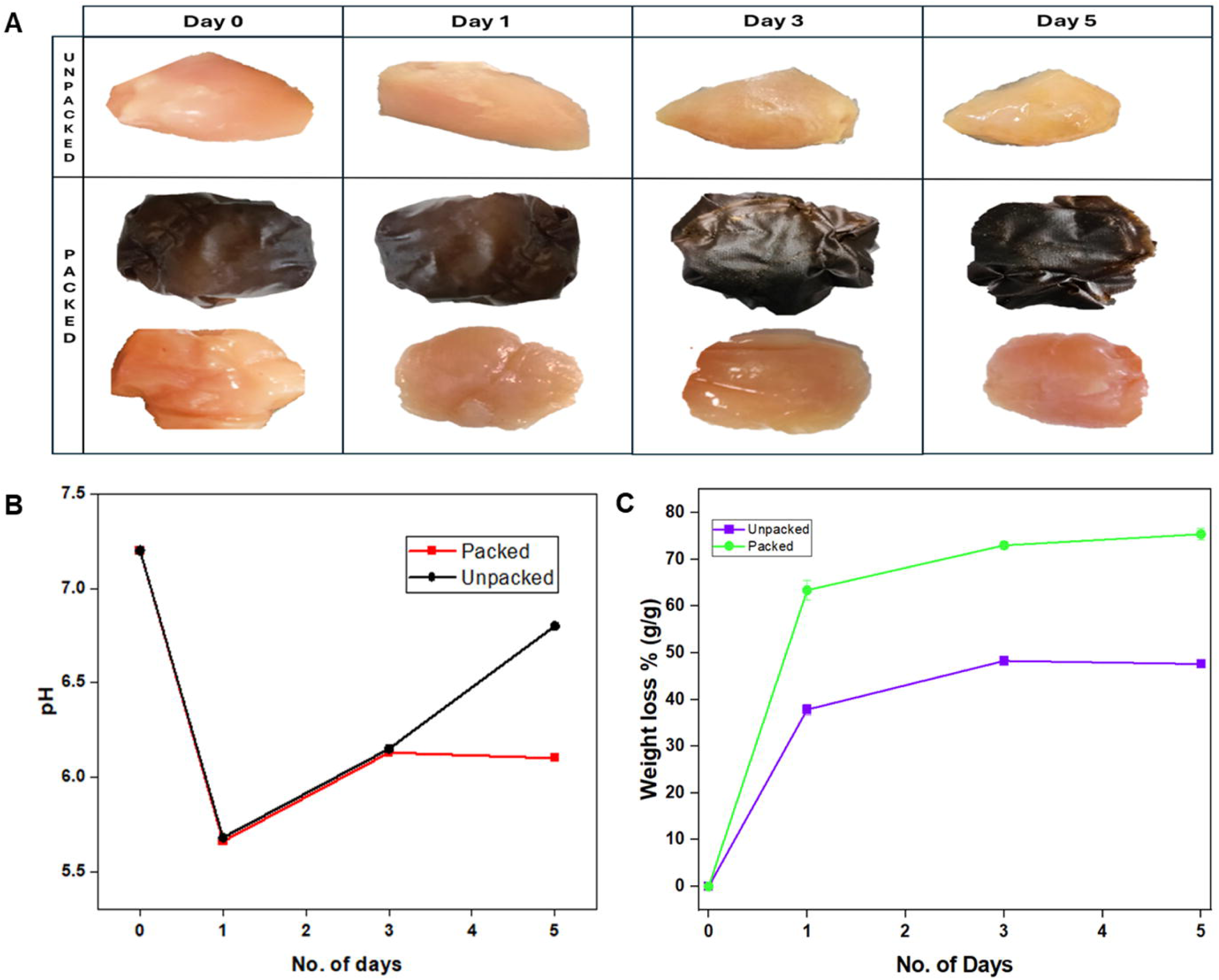
(A) Total microbial count (log[[CFU/mL), and (B) soluble protein content of packed and unpacked chicken breast samples during storage at 25°C.

#### 3.16.4 Soluble protein content

Soluble protein concentration (g/g) in chicken breast was analysed to evaluate protein stability during storage in both the test (packed)and control (unpacked) sample (Fig. 10B). On Day 0, both the chicken samples exhibited soluble protein levels of 0.196 ± 0.009 g/g. By Day 5, the test sample (packed) exhibited substantially lower soluble protein content of 0.50 ± 0.013 g/g, compared to 0.73 ± 0.082 g/g in the unpacked control indicating restricted oxygen exchange within the hydrogel packaging and maintaining protein stability by limiting oxidative and enzymatic degradation. Oxidative reactions can lead to the formation of undesirable compounds and off-odors and adversely affect the texture and water-holding properties of meat through protein breakdown and denaturation(Wang et al., 2022). In contrast, the unpacked sample exhibited an enhanced solubilization of myofibrillar proteins driven by endogenous protease (calpains and cathepsins) and microbial proteases (peptidyl-, amino-, and carboxypeptidases) that promote hydrolysis into peptides and free amino acids (Abril et al., 2023). The lower increment in soluble protein in the packed samples confirms that the hydrogel film effectively suppressed protein denaturation and oxidative reactions, thereby preserving protein integrity and delaying spoilage, while reducing off-odors and textural deterioration.

#### 3.16.5 Malondialdehyde (MDA) analysis

Lipid oxidation is a primary cause of quality deterioration in stored meat products, resulting in rancid flavour, discoloration, and reduced shelf life. MDA is a secondary oxidation product that was used to monitor lipid peroxidation in both test (packed) and control (unpacked) chicken breast samples stored at 25 °C Fig. 11). At Day 0, both groups showed similar baseline MDA levels of 0.58 nmol/g, indicating minimal initial oxidation. By Day 5, the unpacked samples reached 3.62 nmol/g, while those packed with the CL-0.8Lig/3Cht film showed a lower value of 2.30 nmol/g, reflecting around 37% reduction in oxidative load. This attenuation of lipid peroxidation indicates that the bio composite film substantially delays the onset of spoilage, extending the oxidative stability window at 25°C. The suppression effect is attributed to the synergistic roles of lignin and chitosan within the composite matrix. Lignin functions as a peroxyl radical quencher by donating electrons to neutralize reactive species released from meat sample, thereby interrupting the lipid peroxidation chain. Additionally, the chitosan matrix provides a dense polymeric network, reducing oxygen permeability (Cazón and Vázquez, 2020) to the meat surface. In contrast to conventional active packaging systems requiring additive loading, the synthesized composite film derives its activity from the structural integration of functional biopolymers.

**Fig. 11.**
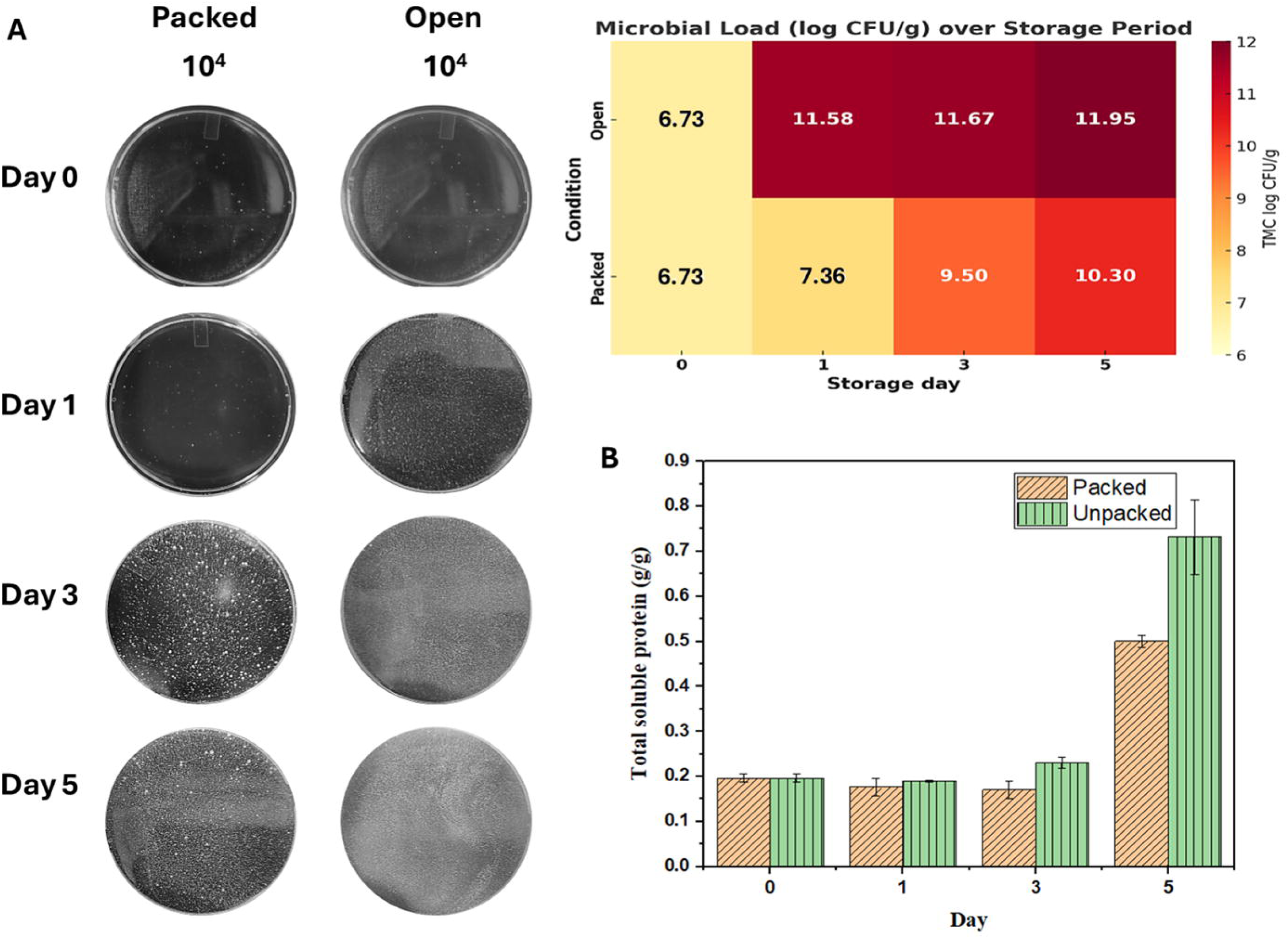
Schematic illustrating CL-0.8Lig/3Cht film mediated reduced lipid peroxidation through ROS scavenging supported by lower MDA levels in packed samples over 5 days.

**Fig. 12.**
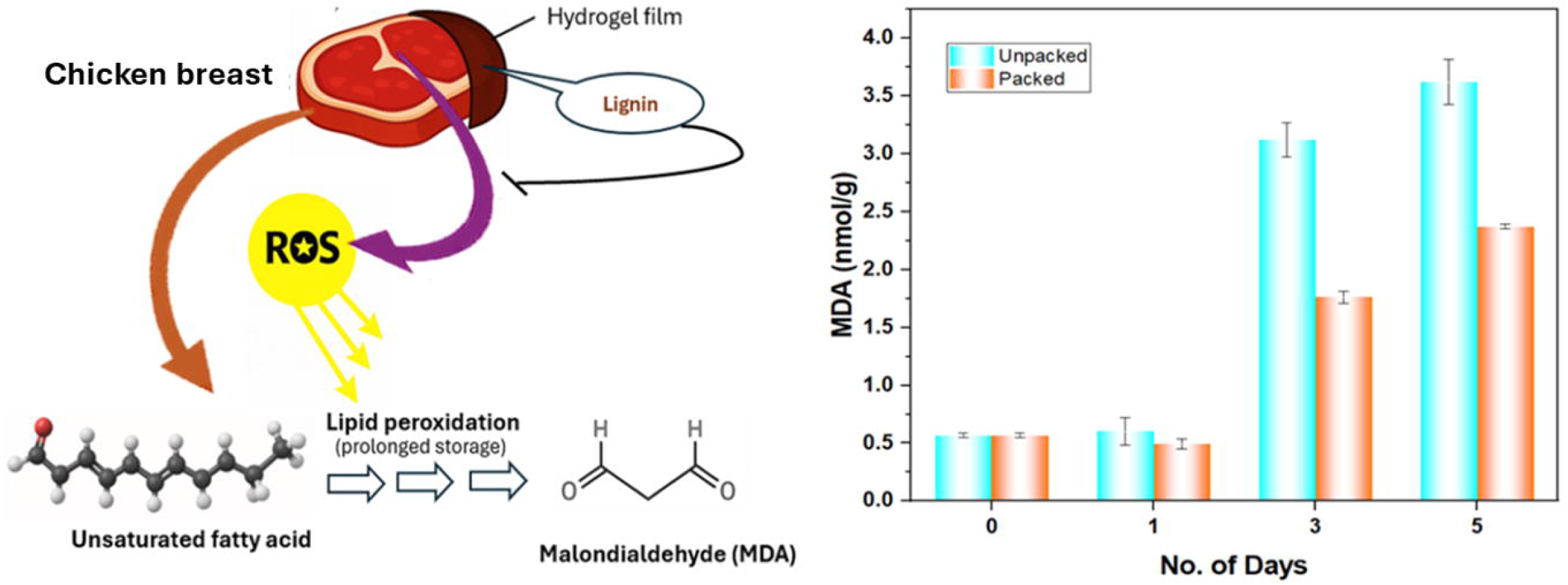
**Biodegradability of 3Cht, 0.8Lig/3Cht, CL-0.8Lig/3Cht films over 90 days.**

### 3.17 Biodegradation study

The biodegradability of the hydrogel films was evaluated through soil burial for 90 days, showing a clear time-dependent degradation trend (Fig. 13). The pure chitosan film (3Cht) degraded rapidly after 40 days, with weight loss increasing from 7.56% on Day 10 to about 95% by Day 60. The 0.8Lig/3Cht exhibited improved resistance, with 3.85% mass loss on Day 10 and 61.94% by Day 90. The presence of lignin slowed down initial microbial attack, possibly due to the aromatic and recalcitrant nature of lignin, but the matrix remained sufficiently biodegradable over time. The CL-0.8Lig/3Cht film showed slower degradation rate, with only 1.98% loss within 20 days and 68.80% by Day 90, confirming that STMP crosslinking significantly delays onset of breakdown, likely by stabilizing the polymer network through phosphate bridging. This feature can make the film structurally stable throughout its utilisation phase that ensures product protection and can also subsequently undergo complete biodegradation after disposal under natural conditions(Cheng et al., 2024).

### 4.0 Conclusions

The present work demonstrates a sustainable cleaner production approach to develop multifunctional hydrogel films for active food packaging through the synergistic integration of chitosan, lignin, and phosphate crosslinking chemistry. The system effectively overcomes the key limitations of conventional biopolymer films such as poor aqueous resistance, the strength–flexibility trade-off and toxic crosslinker utilisation while maintaining biodegradability and material safety. By valorising agricultural lignin along with chitosan and food safe STMP, the study advances circular bioeconomy goals that can reduce reliance on petroleum-based plastics. Furthermore, chicken breasts preservation trials under ambient conditions revealed the film’s potential in extending shelf life by minimizing microbial growth, lipid oxidation, weight loss, and protein denaturation, while maintaining pH stability. Future work may explore scalability, life-cycle assessments, techno-economics, consumer safety, and compatibility with other food types to support industrial translation.

## Acknowledgements

The authors express their gratitude to CSIR-ASPIRE (Grant No. 25WS[56216]/2023) and Anusandhan National Research Foundation (SERB), Ministry of Science and Technology, Govt. of India (Grant No. SRG/BT/F304/2023-24/G686) for providing financial support for this research. They also acknowledge the Council of Scientific and Industrial Research (CSIR), India for awarding the CSIR-NET Junior Research Fellowship (Grant No. 09/1001(15595)/2022-EMR-1). The authors further thank the Director of IIT Hyderabad for enabling this work through access to institutional facilities and support infrastructure.

## Author Contributions

Ms. Sumona Garg: Writing – original draft, data curation, methodology, investigation, and conceptualisation.

Dr. Althuri Avanthi: Visualization, conceptualisation, supervision, project administration; Resources, Writing - review & editing.

---

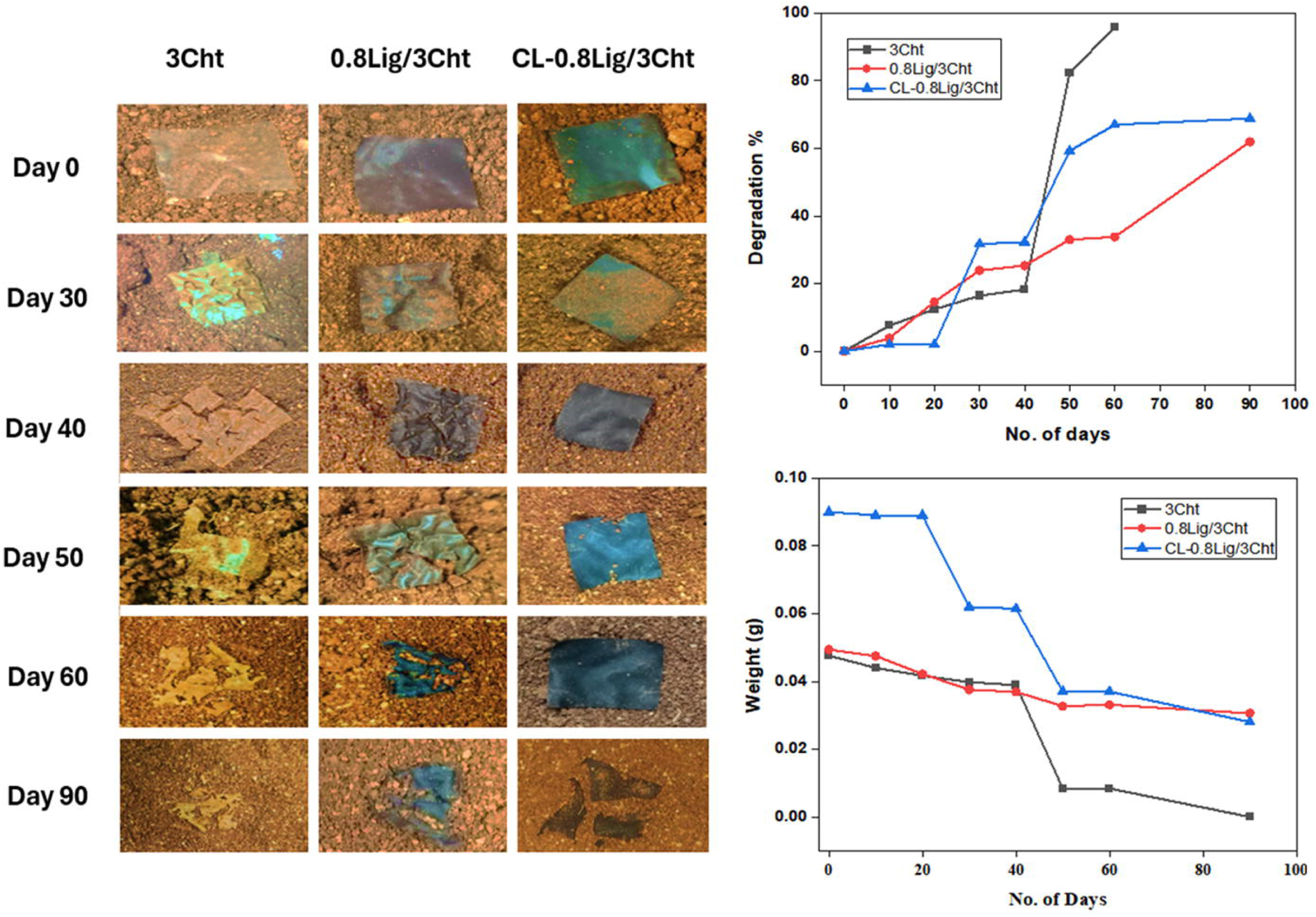

