## Supplemental Fig S1 and Table S1 for "Strong STMP-Crosslinked Lignin/Chitosan Hydrogel Films with Enhanced Aqueous Stability and Bioactivity for Active Food Packaging"

**Supplementary Information**

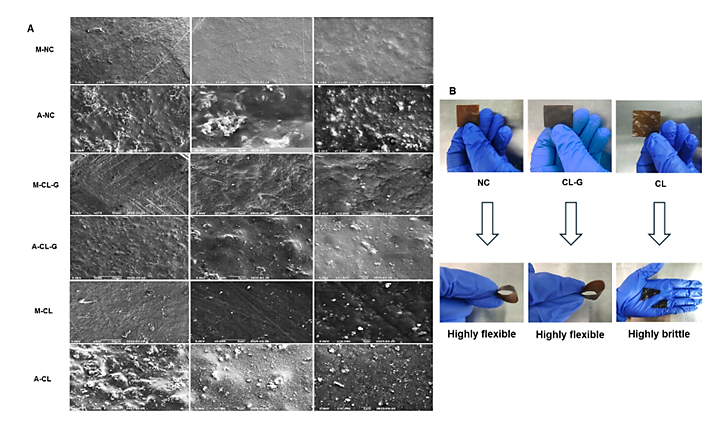

**Fig. S1** (A) M-NC: Matrix side of non-crosslinked 0.8Lig/3Cht film; A-NC: Air side of non-crosslinked film; M-CL-G: Matrix side of crosslinked film with glycerol; A-CL-G: Air side of crosslinked film with glycerol; M-CL: Matrix side of crosslinked film without glycerol; A-CL: Air side of crosslinked film without glycerol, and (B) biocomposite hydrogel film behaviour with and without crosslinking/plasticizer.

**Table S1** UV Absorption characteristics of lignin

| **Wavelength (nm)** | **Type of Transition** | **Lignin Structural Feature** |
| --- | --- | --- |
| **∼310 nm** | π → π* | π → π* in Cα═Cβ linkages conjugated with aromatic ring |
| **∼280 nm** | n → π* | Non-conjugated free or etherified hydroxyl groups |
| **∼240 nm** | n → π* | Non-conjugated hydroxyl groups |
| **200–210 nm** | π → π* | Aromatic ring systems |
